# Bedtime to the brain: How infants sleep habits intertwine with sleep neurophysiology

**DOI:** 10.1101/2021.11.08.467800

**Authors:** Sarah F. Schoch, Valeria Jaramillo, Andjela Markovic, Reto Huber, Malcolm Kohler, Oskar G. Jenni, Caroline Lustenberger, Salome Kurth

**Affiliations:** Department of Pulmonology, University Hospital Zurich, Zurich, CH; Center of Competence Sleep & Health Zurich, University of Zurich, Zurich, CH; Donders Institute for Brain, Cognition and Behaviour, Radboud University Medical Centre, Nijmegen, NL; Child Development Center, University Children’s Hospital Zurich, Zurich, CH; Surrey Sleep Research Centre, Faculty of Health and Medical Sciences, University of Surrey, Guildford, UK; Neuromodulation Laboratory, School of Psychology, University of Surrey, Guildford, UK; Department of Child and Adolescent Psychiatry and Psychotherapy, Psychiatric Hospital, University of Zurich, CH; Children’s Research Center, University Children’s Hospital Zurich, University of Zurich (UZH), Zürich, Switzerland; Neural Control of Movement Lab, Institute of Human Movement Sciences and Sport, Department of Health Sciences and Technology, ETH Zurich, 8092 Zurich, Switzerland; Department of Psychology, University of Fribourg, Fribourg, CH

**Keywords:** EEG, development, infancy, slow wave activity, spindles, coherence, sensitive period, sleep regulation, brain maturation

## Abstract

Adequate sleep is critical for development and facilitates the maturation of the neurophysiological circuitries at the basis of cognitive and behavioral function. Observational research has associated sleep problems in early life with worse later cognitive, psychosocial, and somatic health outcomes. Yet, the extent to which day-to-day sleep habits in early life relate to neurophysiology - acutely and long-term - remains to be explored. Here, we report that sleep habits in 32 healthy 6-month-olds assessed with actimetry are linked to fundamental aspects of their neurophysiology measured with high-density electroencephalography (hdEEG). Our study reveals four key findings: First, daytime sleep habits are linked to EEG slow wave activity (SWA). Second, habits of nighttime movement and awakenings from sleep are connected with spindle density. Third, habitual sleep timing is linked to neurophysiological connectivity quantified as Delta-coherence. And lastly, Delta-coherence at age 6 months predicts nighttime sleep duration at age 12 months. These novel findings widen our understanding that infants’ sleep habits are closely intertwined with three particular levels of neurophysiology: sleep pressure (determined by SWA), the maturation of the thalamocortical system (spindles), and the maturation of cortical connectivity (coherence). Our companion paper complements this insight in the perspective of later developmental outcomes: early thalamocortical connectivity (spindle density) at age 6 months predicts later behavioural status at 12 and 24 months. The crucial next step is to extend this concept to clinical groups to objectively characterize infants’ sleep habits “at risk” that foster later neurodevelopmental problems.

**Highlights:**

- Infant’s habitual sleep behavior (actimetry) is linked with their sleep neurophysiology (EEG)
- Habits of daytime sleeping (naps) are related to slow wave activity
- Infant’s movements and awakenings at nighttime are linked to their sleep spindles
- Sleep timing (infant’s bedtimes) is associated with cortical connectivity in the EEG

## 1. Introduction

Adequate sleep is essential to human health, as it fosters optimal neuronal functioning and cognitive performance (Prince and Abel, 2013; Xie et al., 2013). Sufficient continuous sleep is furthermore crucial for neurodevelopment, as causally demonstrated in animal models (Frank et al., 2001; Shaffery et al., 2002) or correlatively reported in children (Simola et al., 2014). The behavioral state of sleep, allows for the maturation of neurophysiological circuitries, which are considered the basis for cognitive and behavioral functioning. Yet, whether day-to-day sleep habits determine the outcome of brain physiology has not been examined in early human infancy - the most vulnerable developmental period of the lifespan, which also shows the most variable sleep behavior (Iglowstein et al., 2003).

During infancy, sleep habits show the largest individual variability of the human lifetime (Iglowstein et al., 2003). Notably, in approximately 30% of infants, sleep is considered a burden (Lam et al., 2003). Observational research links early sleep behaviors to problematic later psychosocial, cognitive, and somatic outcomes (Gregory et al., 2009, 2005; Mindell et al., 2017; Simola et al., 2014). Thus, clinical reference values are essential for predicting or identifying possible sleep-related risks for subsequent mood disorders or cognitive performance issues later in life. Although neurophysiological features, *e*.*g*., slow waves and spindles in the electroencephalogram (EEG), were proposed to link sleep behavior and brain functioning (Kurth et al., 2012), such systematic examination is still lacking. Particularly in the most vulnerable period for neuronal development - infancy - it remains entirely unexplored how sleep habits are intertwined with neurophysiological features.

In recent years, high-density (hd)EEG has evolved as a new powerful pediatric imaging method as it is non-invasive and provides good spatial and temporal resolution (Lustenberger and Huber, 2012). The increased use of hdEEG led to the discovery that topographical maps of sleep undergo maturation across childhood (Kurth et al., 2010). Analytical advances revealed specific sleep hdEEG features associated with aspects of neuronal function. Consequently, four features from non-rapid eye movement (NREM) sleep have proven to be of specific interest for infant research. The first core EEG feature is slow wave activity (SWA; EEG power between 0.75-4.25 Hz) which indicates the homeostatic regulation of sleep need, *i*.*e*., increasing with the duration of prior wakefulness and decreasing during the time spent asleep (Achermann et al., 1993; Jenni et al., 2005). However, infant homeostatic sleep regulation remains understudied, and contradictory findings exist, *e*.*g*., whether it reflects homeostatic across-night decline (Jenni et al., 2004; Schechtman et al., 1994). The second core feature of infant sleep EEG is theta activity (EEG power in 4.5 - 7.5 Hz), which declines in the course of the night and was therefore proposed to potentially reflect the dissipation of sleep pressure (Jenni et al., 2004). However, which of the two markers - SWA or theta activity - more reliably reflects sleep pressure build-up across wakefulness in infants remains unknown. The third core EEG feature is spindles in the sigma frequency range (11-16 Hz): waxing and waning oscillations generated by the thalamocortical system that may serve a sleep-protective function by preventing waking up from external perturbations during sleep (Andrillon and Kouider, 2020; Fernandez and Lüthi, 2020). Finally, the fourth core EEG feature in infancy is neuronal connectivity, as reflected in EEG coherence. Coherence indicates cortical connectivity by quantifying the EEG synchronicity between electrodes from different locations (Markovic et al., 2020; Tarokh et al., 2010). Cortical connectivity assessed in sleep is a trait-like characteristic property that emerges as a consequence of both internal (genetic, maturational) and external influences (environmental). External influences were identified as the more substantial contributor (Markovic et al., 2020), which triggers the question to what extent parents’ actions, e.g., influencing infants’ sleep timing, can modify cortical connectivity of infant sleep. Importantly, these four core EEG features are also related to learning and plasticity processes (Born and Feld, 2012; Cairney et al., 2018; Holz et al., 2012; Mölle et al., 2004; Tarokh et al., 2014; Wilhelm et al., 2014). For example, SWA indicates episodic memory processing in children’s naps (Lokhandwala and Spencer, 2021) and increased spindle content in naps related to successful learning in a generalization task (Friedrich et al., 2019). Our companion paper demonstrates that sleep spindles in healthy infancy predict their later developmental outcomes (Jaramillo et al., 2021). Yet, whether infants’ sleep habits, *i*.*e*., napping behavior, bedtimes, and nighttime awakenings, connect to the four primary sleep EEG features remains unexplored.

Adults’ sleep habits are manifested in their neurophysiology, as shown by increased SWA in long vs. short sleepers after sleep deprivation (Aeschbach et al., 1996) and more low sigma power for early vs. late chronotypes (Mongrain et al., 2005). Effects of sleep habits on EEG characteristics are also evident in adolescents; morning types, for example, exhibit reduced spindles (Merikanto et al., 2017). Beyond its relationship with neurophysiology, sleep duration and bedtimes in adolescence are linked to brain morphology (gray matter volume) and behavioral performance (Urrila et al., 2017).

While the link between sleep habits and neurophysiology is entirely unexplored in infants and young children, acute deviations from otherwise regular sleep schedules significantly affect their neurophysiology. For example, habitually napping toddlers who miss a nap demonstrate a substantial homeostatic response at nighttime, with the typical increased power in SWA (Lassonde et al., 2016). Similarly, acute sleep deprivation in young children raised sleep pressure and thus triggered a homeostatic response of increased SWA, specifically affecting developing brain areas (Kurth et al., 2016). Therefore, it is conceivable that chronically extending or restricting young children’s sleep beyond their individual needs is a potential risk for neurodevelopment and could affect emotional lability, impulsivity (Gruber et al., 2012), and neurocognitive functioning (Molfese et al., 2013). In addition, it is crucial to consider an infant’s sleep in the complex context it is experiencing daily: while some sleep habits are more “internally” determined in infants (e.g., fragmentation of sleep, total sleep need (Franken et al., 2001)), other infant sleep habits depend on both internal needs with external influences of the parents (e.g., sleep timing). Fortunately, an infant’s internal and external sleep habits can be summarized with five sleep composites (Schoch et al., 2020). These five core domains can be used to capture the complexity and multi-dimensionality of sleep behaviors in the first year of life. They include *Sleep Activity*, reflecting movement and awakenings at night; *Sleep Timing*, summarizing the clock time of bedtimes and sleep times; *Sleep Night*, characterizing nighttime sleep opportunity and duration; *Sleep Day*, representing duration and number of daytime naps and their regularity; and *Sleep Variability*, reflecting the variability of timing and nighttime sleep between the recorded days.

In summary, adults’ and adolescents’ sleep habits are manifested in their neurophysiology, with specific EEG features underlying this link. However, this examination is lacking in infancy, the most vulnerable period of the human lifespan, which sets the foundation for brain and behavioral development. We thus tested the hypothesis that infants’ sleep habits at age 6 months are closely linked to infants’ neurophysiology at 6 months assessed with hdEEG during sleep. To characterize this linkage, we investigated correlations between infant’s SWA, theta power, spindle density, and Delta-coherence with sleep habits, as quantified from actigraphy (core composites *Sleep Day, Sleep Night, Sleep Timing, Sleep Variability*, and *Sleep Activity*) (Schoch et al., 2020). We also evaluated the topographical dimension of these associations across the scalp. Further, we tested whether infants’ sleep behaviors infer sleep neurophysiology or *vice versa*. Longitudinal recordings at ages 3, 6, and 12 months provided the basis for examining the direction of predictive associations.

## 2. Methods

### 2.1 Participants

We invited parents with healthy 6-month-old infants enrolled in a research project tracking infant sleep (Schoch et al., 2021, 2020) to participate in an at-home EEG assessment. Of 152 families, 24 agreed to participate in the EEG recording. Additionally, we recruited 11 families for an hdEEG sleep assessment and sleep tracking at infant age 6 months. Of the 35 participants, 32 (n = 15 female) were included in the final analyses, as three were excluded due to incomplete recordings (n = 2 inability to fall asleep, n = 1 too short sleep period). Infants were healthy (i.e., absence of central nervous system disorders, acute pediatric disorders, brain damage, chronic disease, family background of narcolepsy, psychosis, or bipolar disorder), primarily breastfed (> 50% of feedings), and had no antibiotics intake until age 3 months.

They had been delivered vaginally and at term with a birth weight above 2500 g. Parents were required to have good knowledge of the German language. Ethical approval was obtained from the *cantonal ethics committee* (BASEC 2016-00730), and study procedures adhered to the declaration of Helsinki. Parents gave written informed consent after an explanation of the study. Families received small gifts for their participation, including a 40 CHF voucher.

### 2.2 Experimental design

We used actigraphy at 3, 6, and 12 months in 22 infants or at 6 months only in 10 infants to assess sleep habits. At each assessment, infant sleep habits were measured for approximately 11 continuous days using ankle actigraphy (GENEactiv, Activinsights Ltd, Kimbolton, UK) and parents filled in a concurrent 24-h-diary. Actigraphs were worn continuously and only removed for bathing/swimming, which caregivers documented.

### 2.3 hdEEGassessed at families’ homes

Neurophysiological markers were obtained at 6 months using sleep hdEEG. EEG measurement was conducted at the families’ home using a high-density sponge electrode net and scheduled to each infant’s habitual bedtime (124 electrodes, Electrical Geodesics Sensor Net, Electrical Geodesics Inc., EGI, Eugene, OR). Nighttime sleep was measured for a maximum of 2 hours. The net was soaked in electrolyte water (1 l) containing potassium chloride (1%) and baby shampoo for 5 minutes before application. After applying the electrode net, impedances were kept below 50 kΩ. EEG data were referenced to the vertex during recording, sampled at 500 Hz with a bandpass filter (0.01, 200 Hz).

### 2.4 Quantifying infant sleep habits

According to our laboratory standards based on actigraphy analysis, sleep habits were quantified as five infant sleep composites (Schoch et al., 2020). In brief, we computed sleep-wake patterns at a 1-minute resolution using a 6-step modification (Schoch et al., 2019) of the Sadeh algorithm (Sadeh et al., 1995), taking into account the sleep diaries. Using a principal component analysis, we derived 5 sleep composites from 48 commonly-used sleep variables based on sleep-wake patterns (PCA, (Schoch et al., 2020). The five sleep composites are *Sleep Day, Sleep Night, Sleep Activity, Sleep Timing*, and *Sleep Variability. Sleep Day* includes daytime nap duration, number, and regularity. *Sleep Night* includes nighttime sleep opportunity and duration. *Sleep Activity* includes movements and awakenings at night. *Sleep Timing* characterizes the clock time of bed- and sleep times. *Sleep Variability* summarizes the variability of timing and duration of nighttime sleep between the recorded days. In case of missing a single recording day, missing sleep variables that were part of the PCA were imputed.

### 2.5 hdEEG analysis

EEG analysis was performed with Matlab (R2020b). First, EEG data was bandpass filtered (0.5 - 50 Hz) and down-sampled to 128 Hz. Sleep stages were visually scored by two independent raters according to the AASM Manual (Iber et al., 2007), using 20 s epochs and discussing to reach consensus for final scoring. Artifacts were rejected by visualizing frequency and power for both channels and time, using a semiautomatic approach (Huber et al., 2000). Afterward, the EEG was re-referenced to average reference. The outermost electrodes (Kurth et al., 2010) and additional channels (max. 10 %) with a low percentage of good epochs were removed, resulting in recordings from 74 - 109 electrodes (M = 100.5, SD = 7.5). Missing electrodes (excluded due to artifacts) were interpolated for topographical analysis.

EEG Power was determined for each electrode in the slow wave activity (SWA, 0.75-4.25 Hz) and theta (4.5 - 7.5 Hz) frequency range, averaged across the first 25 - 30 minutes of artifact-free NREM sleep (30 minutes unless less data was available n = 3). In the same data, spindles were detected automatically using an approach similar to (Ferrarelli et al., 2007; Gerstenberg et al., 2020; Lustenberger et al., 2015) as described in the companion paper (Jaramillo et al., 2021). Fast spindle density (13.5 - 16 Hz) was quantified as the average number of spindles detected per minute.

Coherence was calculated between all possible pairs of electrodes using Welch’s method. Coherence values range from 0 to 1 and indicate the level of correlation between two signals at a specific frequency. Specifically, coherence was defined as |P_xy (f)|^2/(P_xx (f)P_yy (f)) where P_xy (f) is the cross-spectral density and P_xx (f) and P_yy (f) are the auto-spectral density functions of the two signals x and y at frequency f (Bendat and Piersol, 2011). To limit the number of electrode pairs, i.e., connections, we applied the data-driven clustering method described by ten Caat et al. (2008) across the SWA, theta, and fast spindle frequency ranges. However, as the results were similar and there was a high correlation among coherence data of the three frequency ranges, we limited our analyses to the SWA frequency range (correlation Delta-coherence and theta-coherence r(29) = 0.81, p < 0.001, Delta-coherence and high sigma coherence r(29) = 0.41, p = 0.02). The clustering method partitioned electrodes into spatially connected regions based on the coherence of each electrode with its neighbors. Coherence between pairs of resulting regions was analyzed as the average coherence across all connections with one electrode belonging to one region and the other electrode belonging to the other region. Between-region connections exceeding a predefined threshold for coherence (i.e., 0.5) were detected.

### 2.6 Statistical analysis

Statistical analysis was performed in R Studio (1.3.959 using R version 4.0.0) with the following packages *mice* (Buuren and Groothuis-Oudshoorn, 2011), *dplyr* (Wickham et al., 2015), *tidyr* (Wickham and Henry, 2019), and *magrittr* (Bache and Wickham, 2014). To examine global effects, we averaged the four EEG features across all electrodes. We used generalized linear models with the EEG features as the outcome and sleep habits at 6 months (5 sleep composites) as predictors while controlling for exact age, sex, and breastfeeding status (yes/no, corrected model). We performed an uncorrected model with only the sleep composite and age as predictors if a predictor reached p < 0.1 in the corrected model. For power and spindles, we additionally examined localized effects for each electrode based on one imputation with a partial Pearson correlation (factor age) for each electrode and with statistical nonparametric mapping (SnPM) cluster correction. In brief, the order of the two correlation variables was shuffled randomly, and a Pearson correlation was calculated for each electrode. The maximal number of neighboring electrodes with an r-value above the critical threshold was determined separately for positive and negative r-values. 5000 permutations were performed to obtain a distribution of maximal cluster sizes for positive and negative r-values, and the threshold was set for both distributions to the 97.5^th^ percentile. Next, we tested whether previous sleep habits predict later EEG measures by predicting EEG measures at 6 months by sleep behavior at 3 months while controlling for sex and age at EEG recording. To examine the predictive effects of EEG features for later sleep habits, we ran models using the four EEG features at 6 months as predictors for sleep habits at age 12 months while controlling for age at EEG recording, sex, breastfeeding status, and sleep habits age 6 months. The significance level was set to p < 0.05.

## 3 Results

### 3.1 Localization of sleep in infants: Topographies of SWA, theta, spindle density, and Delta-coherence

We mapped the four neurophysiological features at 6 months of age across the scalp and visualized the topographical distribution of SWA, theta, and spindles (Fig 1). SWA was observed in occipital regions and theta power in frontocentral areas and occipital areas. Spindles showed the highest density in central and frontal areas. Between-region Delta-coherence exceeded the predefined threshold of 0.5 between frontal and frontotemporal regions.

**Fig 1.**
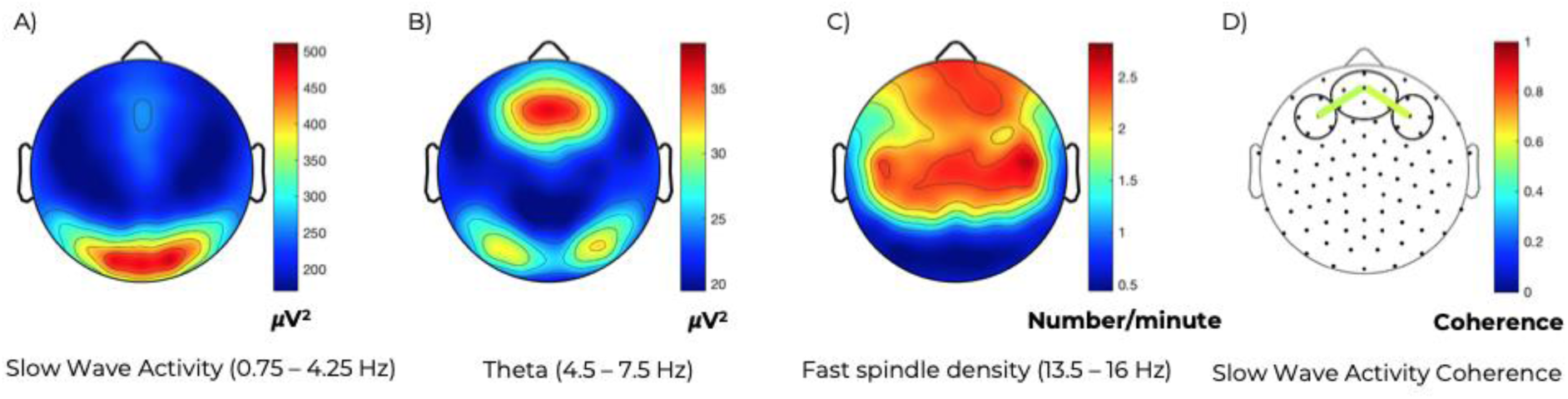
Topographical distribution of the four EEG features across the first 30 minutes of artifact-free NREM sleep at 6 months of age (N = 32). Maps A)-C) are based on 109 electrodes and represent averages across all participants. Values are color-coded and scaled to maximum (red) and minimum (blue). A) Slow wave activity (SWA, 0.75 - 4.25 Hz) is highest in the occipital area. B) Theta power (4.5 - 7.5 Hz) shows the highest presence in frontocentral regions and increased power at occipital regions. C) Fast spindles (13.5 - 16 Hz) show the highest density in central and frontal regions. D) For Delta-coherence, two between-region connections were detected bilaterally between prefrontal and frontotemporal areas (Coherence left = 0.56, Coherence right = 0.58).

### 3.2 Neurophysiological markers are associated with sleep habits at 6 months

Subsequently, we tested whether sleep habits and EEG features at age 6 months are associated. Overall (averaged across all electrodes), daytime sleep habits (*Sleep Day)* were associated with SWA (uncorrected t(25.89) = 2.12, p = 0.04, corrected t(17.98) = 1.85, p = 0.08), i.e., Infants who habitually slept more during the day showed more SWA. Electrode-wise partial correlation between SWA and *Sleep Day* was significant in a large global cluster of 58 electrodes when correcting for the age at the EEG assessment (Fig 2). No other sleep composite was significantly associated with global SWA at 6 months (all p > 0.1, Supplementary Table 1). None of the sleep composites were significantly linked to global theta power (all p > 0.09, Fig 2A).

**Fig 2.**
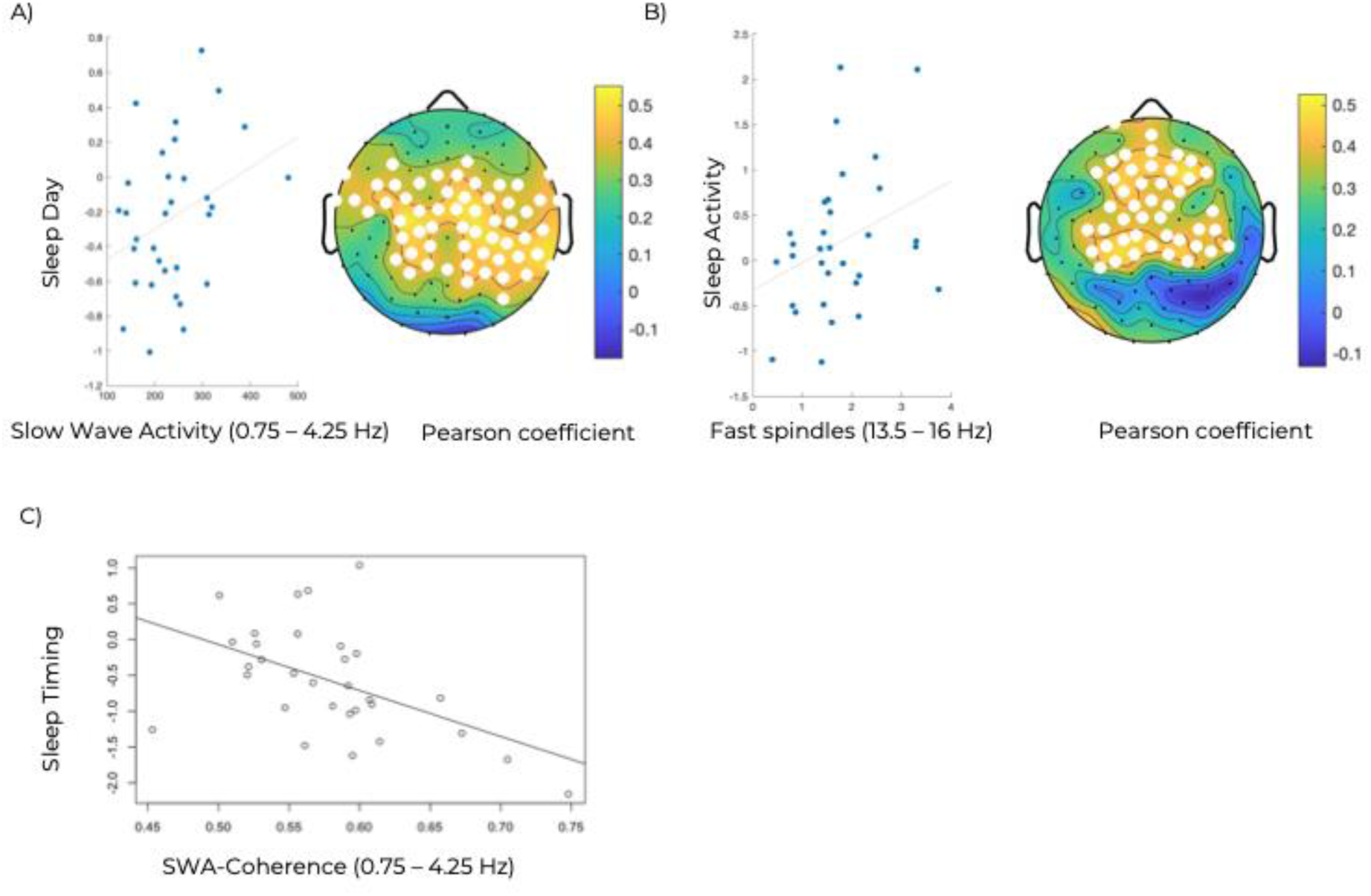
Associations between the 4 EEG features and sleep habits at 6 months with illustrations on a scalp model (N = 32). A) Left: SWA averaged across all 109 electrodes plotted against *Sleep Day (*reflecting daytime sleep habits). Right: Electrode-wise partial correlation between SWA and *Sleep Day* at 6 months of age, corrected for age at EEG assessment. Significant electrodes are indicated as white dots (statistical nonparametric mapping cluster correction for multiple comparisons). B) Left: Fast spindle density averaged across all 109 electrodes plotted against *Sleep Activity* (reflecting movements and awakenings during sleep). Right: Electrode-wise partial correlation between spindle density and *Sleep Activity*, corrected for age at EEG assessment. Significant electrodes are indicated as white dots (statistical nonparametric mapping cluster correction for multiple comparisons). C) Delta-coherence between prefrontal and frontotemporal areas plotted against *Sleep Timing* (reflecting sleep- and bed-times).

In contrast, global spindle density was positively associated with *Sleep Activity* (uncorrected t(26.33) = 2.41, p = 0.02, corrected t(19.26) = 2.27, p = 0.03). Infants whose actimetry revealed more activity and awakenings habitually from sleep showed increased spindle density during the night of the EEG assessment. The electrode-wise partial correlation between spindle density and *Sleep Activity* was significant in a large central cluster of 45 electrodes when correcting for the age at the EEG assessment (p97.5 threshold = 26, Fig 2B). No other sleep composite was significantly associated with global spindle density (all p > 0.1, Supplementary Table 1).

Sleep habits were associated with Delta-coherence at 6 months: *Sleep Timing* was negatively associated with Delta-coherence (uncorrected t(26.01) = -3.11, p = 0.004, corrected t(19.02) = -3.24, p = 0.004, Fig 2C). Infants with earlier sleep timings showed increased Delta-coherence. No other sleep composite was significantly associated with Delta-coherence (all p > 0.05, Supplementary Table 1).

### 3.3 Sleep habits at 3 months do not predict sleep EEG features at 6 months

Next, we explored if sleep behaviors influence long-term neurophysiology. None of the sleep habits at age 3 months consistently predicted EEG features at 6 months (Supplementary Table 2). We also explored if sleep neurophysiology at 6 months predicted infant sleep habits at 12 months. Delta-coherence at 6 months significantly predicted Sleep Night at 12 months (uncorrected t(16.27) = 2.73, p = 0.01, corrected t(10.84) = 1.84, p = 0.09, Supplementary Table 3). Thus, infants with higher Delta-coherence at 6 months had more nighttime sleep at 12 months, indicating that more mature neurophysiology at 6 months predicts more mature sleep behavior at 12 months. SWA, theta, and spindle density were not predictive of later sleep habits.

## 4. Discussion

Our present work on nighttime hdEEG neurophysiology and behavioral sleep habits (as measured by ankle actimetry and sleep diaries) in healthy 6-month-old infants reveals four key findings. First, more pronounced daytime sleep habits are linked to increased SWA across a sizable, globally distributed electrode cluster. Second, more nocturnal activity and awakenings from night sleep are connected with more evident sleep spindles. Third, sleep timing is associated with EEG coherence at age 6 months, and fourth, coherence at age 6 months predicts the duration of nighttime sleep at 12 months. These novel findings widen our understanding that daytime sleep behavior, movement activity at night, and bedtimes of infants are closely intertwined with sleep on the neurophysiological level, which implies that the EEG may be used to monitor whether sleep habits are within norms. The concept that observed sleep behavior is connected to measured sleep in the brain is evident on three distinct levels: sleep pressure (reflecting SWA), the maturation of the thalamocortical system (spindles), and maturation of cortical connectivity (coherence). Considering that sleep habits in early life relate to later maturational outcomes (Spruyt et al., 2008; Timofeev et al., 2020), our study introduces two novel findings: neurophysiology is intertwined with internal and external factors, and infant sleep habits may affect brain development pathways in the long term.

The first main result aligns with the concept that SWA is tightly linked to sleep-wake regulation (Schechtman et al., 1994). Our findings contribute to the insight that infants with increased daytime sleep pressure (*i*.*e*., more pronounced napping) experience nearly globally elevated SWA at night. This finding may seem counter-intuitive considering that napping reduces sleep pressure (and thus SWA) in children (Lassonde et al., 2016) and adults (Werth et al., 1996). However, it can be explained by the specific dynamics of sleep pressure accumulation in infants, *i*.*e*., infants with a fast build-up of sleep pressure might generally experience increased sleep need, both during the day and at night.

Contrastingly, we found no link between infants’ sleep habits and theta power. Interestingly, the dissipation of sleep pressure across sleep in infancy has been suggested to be reflected in theta instead of SWA (Jenni et al., 2004). However, a follow-up study analyzed the same dataset and demonstrated an across-night decrease of the slope of slow waves (Fattinger et al., 2014). Though we only report results concerning SWA, the slope (slope 55, corrected for slow wave amplitude, as in Fattinger et al.) was highly correlated to SWA in our exploratory analyses, potentially indicating that the slope reflects both the sleep pressure build-up as well as its dissipation.

From a spatial dimension, SWA predominated over occipital regions at age 6 months, in line with findings at age 3 months (Guyer et al., 2019) and at preschool age (Kurth et al., 2010). Therefore, we conclude that the localization of SWA on the scalp remains stable across the first years of life. Interestingly, the association between an infant’s daytime sleep habits and SWA is represented in a sizable near-global cluster across central and temporal scalp regions and not in the region of maximal SWA.

As expected, spindle density was maximal over frontal and central scalp areas (D’Atri et al., 2018). The association of spindles with nighttime movements and awakenings aligned with the location of maximal spindle detection. To our knowledge, this is the first demonstration of a link between infants’ nighttime movements/awakenings (*i*.*e*., *Sleep Activity*) with spindle density. This discovery was initially surprising, considering that previous research with human adults reported a “protective role” of spindles for sleep - thereby preventing awakenings (Dang-Vu et al., 2010; Fernandez and Lüthi, 2020; Schabus et al., 2012). However, we acknowledge that *Sleep Activity* computation is based on nighttime actimetric movement. Infants’ recorded nighttime movements may also entail myoclonic twitches, which are the spontaneous movements typically observed in Rapid Eye Movement (REM) sleep (Sokoloff et al., 2020). These twitches provide topographically precise activity to integrate peripheral fiber innervation to the developing central nervous system, as extensively studied in rats (Blumberg et al., 2013; Tiriac et al., 2015). Twitching is thus assumed critical for the microcircuitry development of the sensorimotor cortex. Compelling novel research in human infants discovered that twitches also occur in NREM sleep and in synchrony with sleep spindles (Sokoloff et al., 2021). Thus, our data fundamentally extend the known concept that spindles reflect thalamocortical network strength (Fernandez and Lüthi, 2020) and propose a core linkage of spindles with sensorimotor microcircuitry development in infants. In other words, thalamocortical connectivity underlying infants’ spindles may be enhanced through twitching activity in sleep. The concept that spindles are valuable biomarkers for neurodevelopment is supported by the discovery in our companion paper where we report that spindles at 6 months predict gross motor development at both 12 and 24 months of age (Jaramillo et al., 2021). Our findings highlight the importance of characterizing neurophysiological dynamics across development, as their functions differ according to developmental stage (Purcell et al., 2017).

The third novel finding was that habitual *Sleep Timing* of healthy 6-month-olds was manifested in cortical connectivity (Delta-coherence), such that earlier bedtimes indicated increased cortical connectivity. In a more global context of sleep-wake pattern maturation, the general trend is that sleep increasingly consolidates towards the nighttime (Iglowstein et al., 2003). Thus, earlier infant bedtimes generally signify a more mature pattern. Our novel data confirm this on the neurophysiological level: more mature *Sleep Timing* aligns with more mature cortical connectivity, *i*.*e*., increased connectivity (Kurth et al., 2013; Tarokh et al., 2010). Thus, the effective implementation of infants’ bedtimes should be considered based on the brain’s maturational status.

The spatial dimension also supports this concept, as topographical coherence reached significance only between frontal and frontotemporal areas. Knowing that preterm infants maintain reduced connectivity in exactly frontal regions (Gozdas et al., 2018), our finding suggests that frontal connectivity reflects neurophysiological maturational status.

Furthermore, cortical connectivity is determined by both internal (i.e., demands of the infant’s sleep need) as well as external (i.e., contextual demands of the parents) influences (Markovic et al., 2020). Interestingly, among the five sleep composites (*Sleep Day, Sleep Night, Sleep Timing, Sleep Activity*, and *Sleep Variability*), *Sleep Timing* is likely most reflective of these two. We thus conclude that both sleep timing and cortical connectivity indicate maturational status and represent the multidimensional nature-and-nurture dynamics.

Overall, the main findings of this paper were observed in infants aged 6 months. In addition, we found that Delta-coherence at 6 months predicted nighttime sleep duration at 12 months, indicating that more mature neurophysiology at 6 months (Tarokh et al., 2010) predicts more mature sleep habits at 12 months. This finding could be indicative that the maturing of neurophysiology precedes the development of sleep habits. However, additional associations between infants’ EEG features at age 6 months with either their previous (at age 3 months) or their later sleep habits (at age 12 months) could not be observed. A potential explanation could be the high variability of sleep habits across the first year of life, even within the same infant (Schoch et al., 2020).

Epidemiological research reports the connection between chronically “poor” sleep habits in early life and problematic later psychosocial, cognitive, and somatic outcomes (Lam et al., 2003; Simola et al., 2014). Surprisingly, so far, the underlying neurophysiological links have not been investigated. We thus applied quantitative methodologies to address a fundamental knowledge gap and demonstrated that three sleep EEG features relate to infants’ sleep habits. Interestingly, diverse aspects of sleep habits are associated with distinct features of sleep neurophysiology, illustrating that further advances in our research field require a comprehensive assessment of both - sleep habits and sleep neurophysiology. Furthermore, our results introduce a new concept of the multi-dimensionality of sleep: While sleep habits are known to be intertwined with sleep pressure, influenced by circadian and homeostatic sleep regulators, we show for the first time that sleep habits also closely connect with neuronal connectivity. Consequently, this work highlights the interesting novel translational perspective that sleep habits, which can be modified (*e*.*g*., bedtimes), are connected to neurophysiological maturation processes, which has been beyond our range of influence. Therefore, aligning neurophysiological needs with the behavioral implementation in sleep-wake structuring is essential, both for the infant’s developmental outcome (Mindell et al., 2017; Spruyt et al., 2008) as well as family dynamics and well-being in the long term (Lam et al., 2003).

We assessed infant sleep habits comprehensively at 3, 6, and 12 months of age and conducted a nighttime sleep neurophysiology assessment at the single time point at age 6 months. Repeated measures of sleep neurophysiology would be needed to establish the stability of neurophysiological measures and extrapolate the reported findings to other age groups. They would allow studying the directionality of the relationships by using, for instance structural equation modeling. There are other aspects of the sleep EEG, which we did not include in this study, because of strong correlations with the included variables such as slow spindles, sigma power etc. Future studies will need to investigate potential associations of these variables and sleep habits. Further, experimental alterations of sleep habits (Werner et al., 2015) and sleep neurophysiology (Ngo et al., 2013) are required to establish the directionality of their longitudinal associations.

In conclusion, we report that sleep habits are linked to multiple sleep neurophysiology features in healthy infants. The next crucial step is to characterize which sleep habits represent a risk profile for neurodevelopmental problems to identify infants who benefit most from sleep interventions.

## Acknowledgments & Funding

We thank the parents and infants for participating in our study.

This research was funded by the University of Zurich (Clinical Research Priority Program “Sleep and Health”, Forschungskredit FK-18-047, Faculty of Medicine), the Swiss National Science Foundation (SNSF, PCEFP1-181279, P0ZHP1-178697, PZ00P3_179795, P2ZHP1_195248), Foundation for Research in Science and the Humanities (STWF-17-008), and the Olga Mayenfisch Stiftung.

## CRediT roles

**Sarah F. Schoch**: Conceptualization; Data curation; Formal analysis; Funding acquisition; Investigation; Methodology; Project administration; Software; Validation; Visualization; Writing - original draft; Writing - review & editing. **Valeria Jaramillo**: Conceptualization; Formal analysis; Methodology; Software; Validation; Visualization; Writing - original draft; Writing - review & editing. **Andjela Markovic:** Formal analysis; Methodology; Software; Visualization; Writing - review & editing. **Reto Huber:** Conceptualization; Methodology; Resources; Software; Supervision; Writing - review & editing. **Malcolm Kohler:** Conceptualization; Resources; Writing - review & editing. **Oskar Jenni:** Conceptualization; Methodology; Writing - review & editing. **Caroline Lustenberger:** Conceptualization; Methodology; Supervision; Writing - review & editing. **Salome Kurth:** Conceptualization; Formal analysis; Funding acquisition; Investigation; Methodology; Software; Writing - original draft; Writing - review & editing.

## Data and code availability statements

Data and code are available upon request to the authors, pending ethical approval, and in alignment with consenting framework. R Analysis scripts are available on OSF.

---

**Supplementary Table 1.** Associations between sleep habits and EEG features at 6 months.

|  | Corrected Model |  | Uncorrected Model |  |
| --- | --- | --- | --- | --- |
| SWA | Estimate + SE [95 % CI] | P Value | Estimate + SE [95 % CI] | P Value |
| (Intercept) | -124.2 ± 181.66 [-503.78, 255.38] | 0.50 | -82.88 ± 159.98 [-411.18, 245.42] | 0.61 |
| Exact age | 55.11 ± 30.43 [-8.41, 118.62] | 0.09 | 57.28 ± 27.25 [1.36, 113.19] | 0.05 |
| Female sex | -18.64 ± 29.68 [-80.57, 43.28] | 0.54 |  |  |
| Sleep Day | 71.45 ± 38.66 [-9.78, 152.69] | 0.08 | 69.76 ± 32.86 [2.2, 137.33] | 0.04 |
| Sleep Night | 12.58 ± 15.26 [-19.32, 44.48] | 0.42 |  |  |
| Sleep Activity | 15.27 ± 20.74 [-28.18, 58.72] | 0.47 |  |  |
| Sleep Timing | 0.23 ± 19.83 [-41.16, 41.62] | 0.99 |  |  |
| Sleep Variability | -22.95 ± 20.15 [-65.02, 19.12] | 0.27 |  |  |
| Breastfeeding | 53.74 ± 51.79 [-57.65, 165.14] | 0.32 |  |  |
| **Theta** | **Estimate + SE [95 % CI]** | **P Value** | **Estimate + SE [95 % CI]** | **P Value** |
| (Intercept) | 17.65 ± 19.52 [-23.06, 58.36] | 0.38 | 20.67 ± 18.4 [-17.07, 58.41] | 0.27 |
| Exact age | 2.37 ± 3.28 [-4.46, 9.21] | 0.48 | 0.85 ± 3.09 [-5.49, 7.2] | 0.78 |
| Female sex | -6.38 ± 3.18 [-13.01, 0.24] | 0.06 |  |  |
| Sleep Day | 0.21 ± 3.99 [-8.1, 8.52] | 0.96 |  |  |
| Sleep Night | 0.3 ± 1.62 [-3.07, 3.67] | 0.85 |  |  |
| Sleep Activity | 0.12 ± 2.19 [-4.45, 4.69] | 0.96 |  |  |
| Sleep Timing | 3.74 ± 2.13 [-0.7, 8.18] | 0.09 | 2.31 ± 2.02 [-1.84, 6.47] | 0.26 |
| Sleep Variability | -3.13 ± 2.15 [-7.61, 1.36] | 0.16 |  |  |
| Breastfeeding | -4.14 ± 5.24 [-15.25, 6.97] | 0.44 |  |  |
| **Spindles** | **Estimate + SE [95 % CI]** | **P Value** | **Estimate + SE [95 % CI]** | **P Value** |
| (Intercept) | 3.04 ± 1.85 [-0.84, 6.91] | 0.12 | 4.4 ± 1.8 [0.71, 8.09] | 0.02 |
| Exact age | -0.27 ± 0.31 [-0.91, 0.38] | 0.40 | -0.47 ± 0.31 [-1.09, 0.16] | 0.14 |
| Female sex | -0.55 ± 0.3 [-1.18, 0.08] | 0.08 |  |  |
| Sleep Day | 0.06 ± 0.4 [-0.77, 0.89] | 0.88 |  |  |
| Sleep Night | 0.02 ± 0.16 [-0.31, 0.34] | 0.92 |  |  |
| *Sleep Activity* | *0.47 ± 0.21 [0.04, 0.91]* | *0.03* | *0.47 ± 0.2 [0.07, 0.87]* | *0.02* |
| Sleep Timing | -0.35 ± 0.2 [-0.77, 0.08] | 0.11 |  |  |
| Sleep Variability | 0.22 ± 0.21 [-0.22, 0.66] | 0.31 |  |  |
| Breastfeeding | 0.41 ± 0.56 [-0.82, 1.65] | 0.48 |  |  |

| Coherence | Estimate + SE [95 % CI] | P Value | Estimate + SE [95 % CI] | P Value |
| --- | --- | --- | --- | --- |
| (Intercept) | 0.57 ± 0.13 [0.29, 0.84] | < 0.001 | 0.5 ± 0.12 [0.26, 0.75] | < 0.001 |
| Exact age | 0 ± 0.02 [-0.05, 0.04] | 0.82 | 0.01 ± 0.02 [-0.03, 0.05] | 0.66 |
| Female sex | 0.05 ± 0.02 [0.01, 0.1] | 0.02 |  |  |
| Sleep Day | $0 \pm 0.03 [-0.06, 0.05]$ | 0.87 | | |
| Sleep Night | **$0.02 \pm 0.01 [0, 0.04]$** | **0.05** | $0.02 \pm 0.01 [0, 0.04]$ | 0.12 |
| Sleep Activity | $-0.01 \pm 0.01 [-0.04, 0.02]$ | 0.44 | | |
| *Sleep Timing* | $-0.04 \pm 0.01 [-0.07, -0.02]$ | 0.004 | $-0.04 \pm 0.01 [-0.07, -0.01]$ | 0.004 |
| Sleep Variability | $0.01 \pm 0.01 [-0.01, 0.04]$ | 0.29 | | |
| Breastfeeding | $-0.01 \pm 0.04 [-0.1, 0.08]$ | 0.81 | | |

**Supplementary Table 2.**
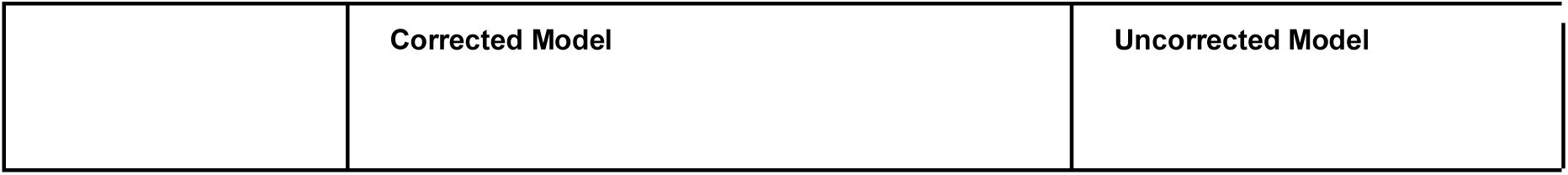
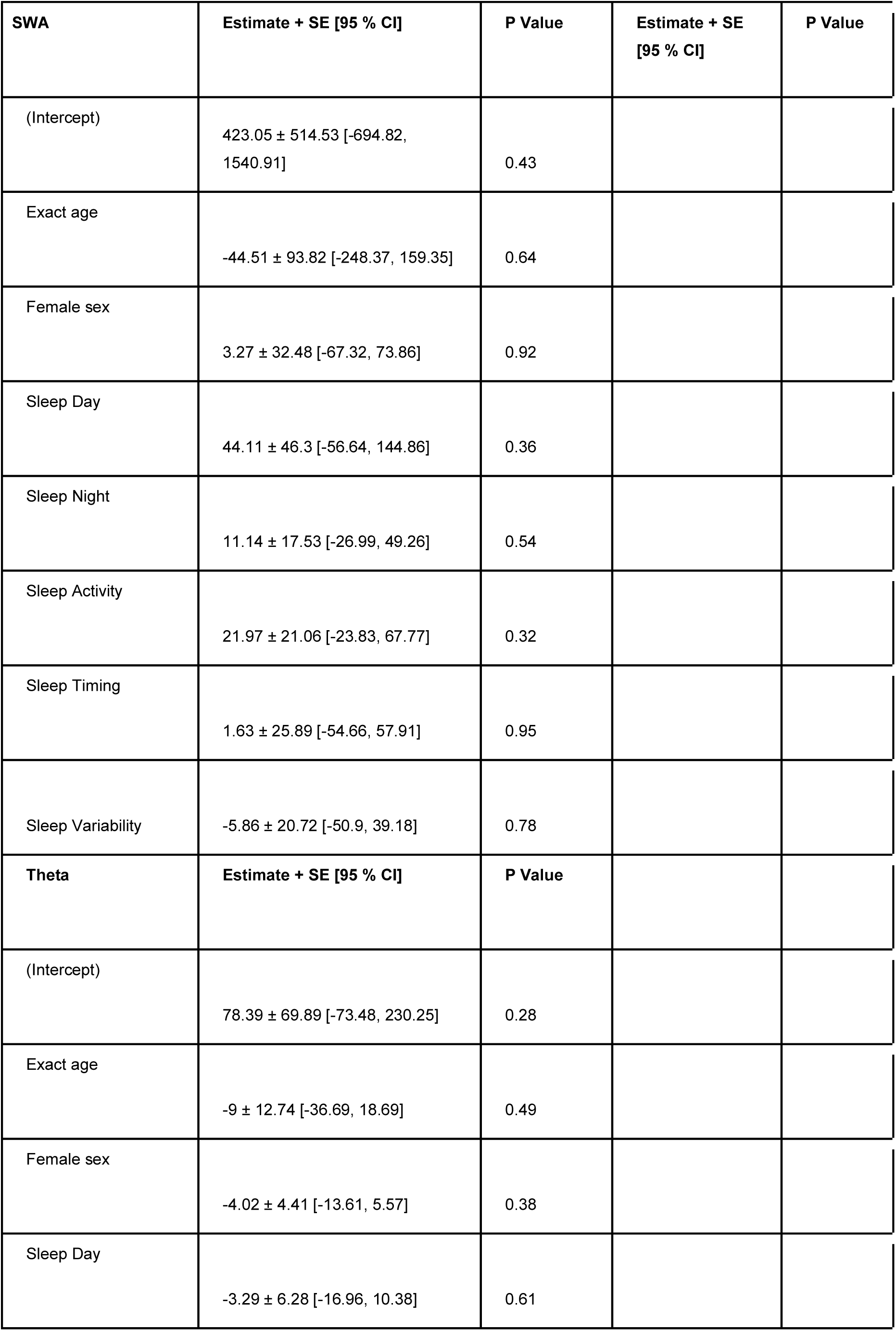
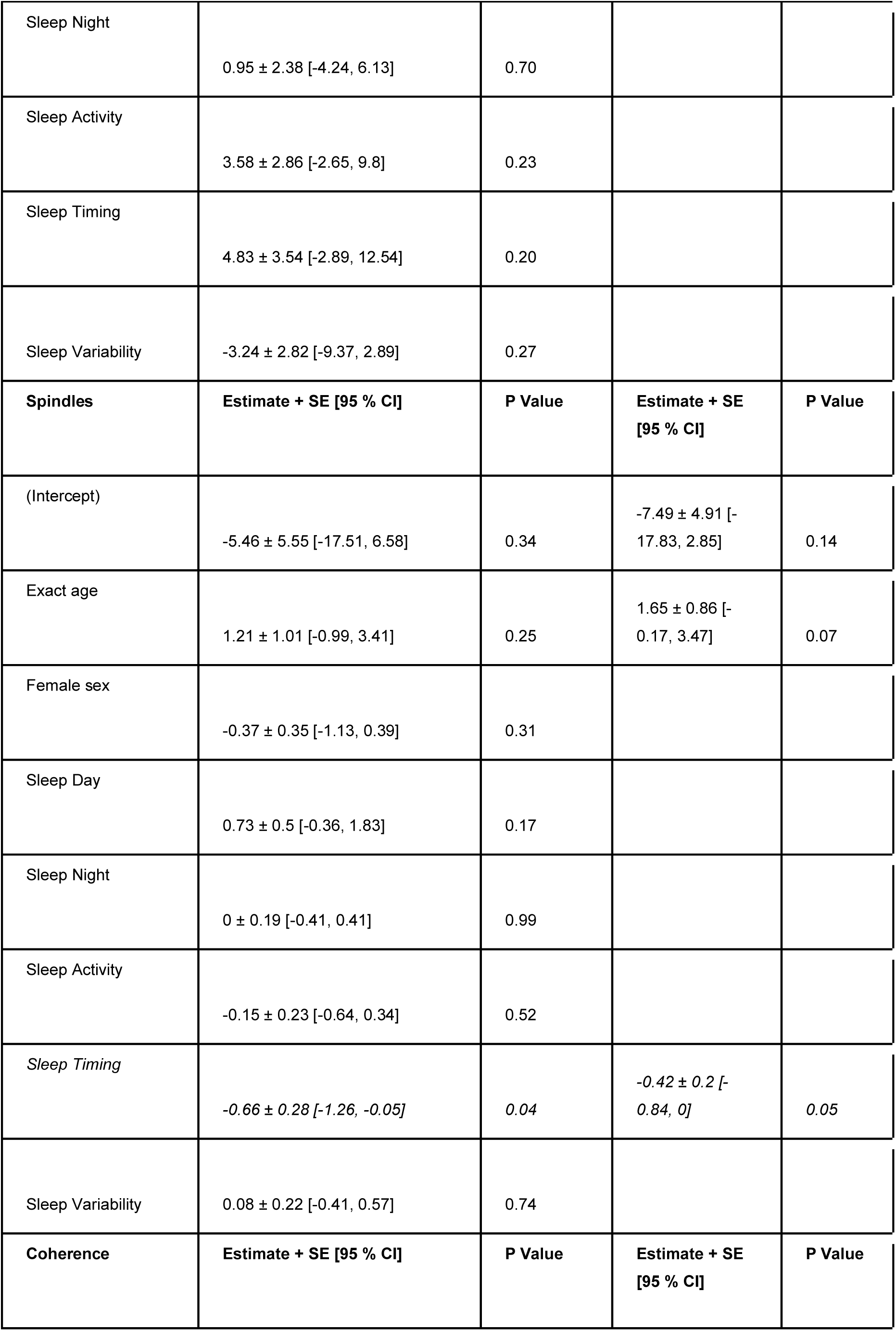
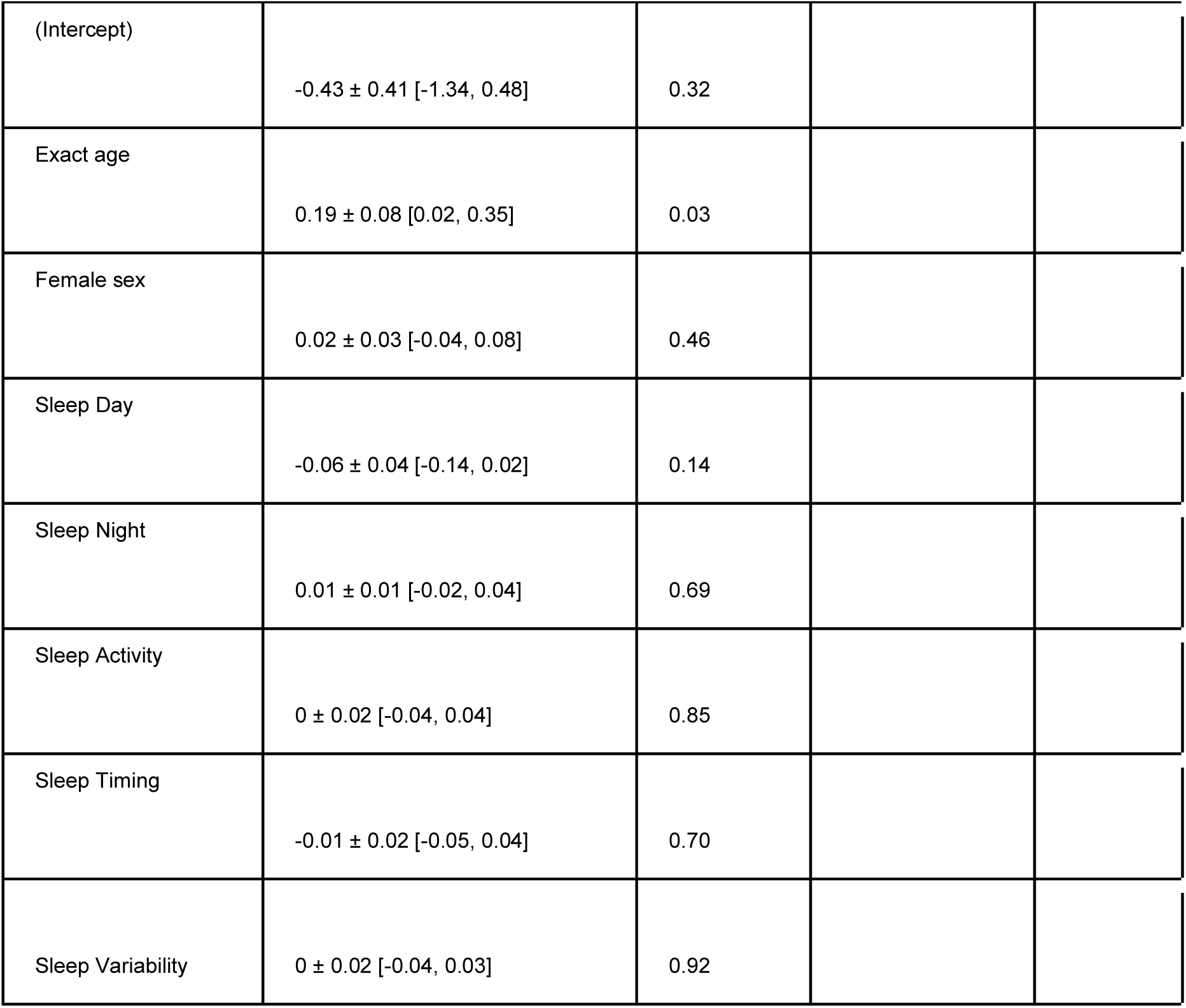
Associations between sleep habits at infants age 3 months and EEG Features at age 6 months.

**Supplementary Table 3.** Associations between EEG features at infant age 6 months and sleep habits at age 12 months.

|  | Corrected Model |  | Uncorrected Model |  |
| --- | --- | --- | --- | --- |
| Sleep Day | Estimate + SE [95 % CI] | P Value | Estimate + SE [95 % CI] | P Value |
| (Intercept) | -4.6 ± 5.3 [-16.26, 7.07] | 0.40 |  |  |
| Exact Age | 0.96 ± 0.97 [-1.17, 3.09] | 0.34 |  |  |
| Female sex | -0.33 ± 0.36 [-1.12, 0.46] | 0.37 |  |  |
| SWA | 0 ± 0 [-0.01, 0.01] | 0.92 |  |  |
| Theta | -0.01 ± 0.02 [-0.06, 0.03] | 0.59 |  |  |
| Spindles | -0.28 ± 0.23 [-0.78, 0.22] | 0.24 |  |  |
| Coherence | -1.39 ± 3.12 [-8.23, 5.45] | 0.66 |  |  |
| Sleep Day 6 Mo | 0.19 ± 0.5 [-0.92, 1.3] | 0.71 |  |  |
| **Sleep Night** | **Estimate + SE [95 % CI]** | **P Value** | **Estimate + SE [95 % CI]** | **P Value** |
| (Intercept) | 1.72 ± 5.98 [-11.51, 14.94] | 0.78 | 7.49 ± 5.62 [-4.43, 19.41] | 0.20 |
| Exact Age | -1.35 ± 1.21 [-4.02, 1.33] | 0.29 | -2.17 ± 1.06 [-4.41, 0.06] | 0.06 |
| Female sex | 0.64 ± 0.35 [-0.14, 1.41] | 0.10 |  |  |
| SWA | 0 ± 0 [-0.01, 0] | 0.70 |  |  |
| Theta | 0.06 ± 0.02 [0.02, 0.1] | 0.01 | 0.03 ± 0.02 [-0.02, 0.08] | 0.2 |
| Spindles | 0.51 ± 0.21 [0.04, 0.99] | 0.04 | 0.22 ± 0.27 [-0.35, 0.79] | 0.42 |
| Coherence | 6.88 ± 3.73 [-1.35, 15.11] | 0.09 | 9.31 ± 3.41 [2.09, 16.53] | 0.01 |
| Sleep Night 6 Mo | 0.25 ± 0.23 [-0.26, 0.76] | 0.31 |  |  |

| **Sleep Timing** | **Estimate + SE [95 % CI]** | **P Value** | **Estimate + SE [95 % CI]** | **P Value** |
| --- | --- | --- | --- | --- |
| (Intercept) | -0.85 ± 3.87 [-9.39, 7.68] | 0.83 | 0.87 ± 4.28 [-8.17, 9.91] | 0.84 |
| Exact Age | 0.26 ± 0.68 [-1.25, 1.77] | 0.71 | -0.08 ± 0.75 [-1.65, 1.49] | 0.92 |
| Female sex | -0.17 ± 0.25 [-0.71, 0.37] | 0.50 |  |  |
| SWA | **0 ± 0 [-0.01, 0]** | **0.06** | **0 ± 0 [-0.01, 0]** | **0.09** |
| Theta | -0.01 ± 0.01 [-0.05, 0.02] | 0.42 |  |  |
| Spindles | -0.01 ± 0.17 [-0.38, 0.36] | 0.94 |  |  |
| Coherence | 0.96 ± 3.17 [-6.13, 8.05] | 0.77 |  |  |
| Sleep Timing 6 Mo | 0.69 ± 0.27 [0.08, 1.3] | 0.03 |  |  |
| **Sleep Activity** | **Estimate + SE [95 % CI]** | **P Value** | **Estimate + SE [95 % CI]** | **P Value** |
| (Intercept) | -11.6 ± 5.19 [-23.02, -0.18] | 0.05 | -10.3 ± 5.14 [-21.13, 0.53] | 0.06 |
| Exact Age | 2.75 ± 0.96 [0.65, 4.86] | 0.02 | 1.77 ± 0.92 [-0.17, 3.7] | 0.07 |
| Female sex | -0.14 ± 0.33 [-0.86, 0.58] | 0.68 |  |  |
| SWA | 0 ± 0 [-0.01, 0.01] | 0.92 |  |  |
| Theta | -0.03 ± 0.02 [-0.07, 0.01] | 0.16 |  |  |
| Spindles | $-0.71 \pm 0.25 [-1.27, -0.14]$ | 0.02 | $-0.33 \pm 0.22 [-0.78, 0.13]$ | 0.15 |
| Coherence | $-5.25 \pm 2.85 [-11.5, 0.99]$ | 0.09 | $-4.09 \pm 3.21 [-10.88, 2.71]$ | 0.22 |
| Sleep Activity 6 Mo | $0.58 \pm 0.27 [-0.03, 1.19]$ | 0.06 | | |
| **Sleep Variability** | **Estimate + SE [95 % CI]** | **P Value** | **Estimate + SE [95 % CI]** | **P Value** |
| (Intercept) | $-12.83 \pm 6.5 [-27.14, 1.48]$ | 0.07 | $-16.24 \pm 5.41 [-27.64, -4.83]$ | 0.01 |
| Exact Age | $2.51 \pm 1.2 [-0.12, 5.15]$ | 0.06 | $2.86 \pm 0.97 [0.82, 4.89]$ | 0.01 |
| Female sex | $-0.25 \pm 0.42 [-1.18, 0.67]$ | 0.56 | | |
| SWA | $0 \pm 0 [-0.01, 0.01]$ | 0.66 | | |
| Theta | $0 \pm 0.02 [-0.06, 0.05]$ | 0.95 | | |
| Spindles | $-0.56 \pm 0.28 [-1.16, 0.05]$ | 0.07 | $-0.34 \pm 0.23 [-0.83, 0.16]$ | 0.17 |
| Coherence | $-0.75 \pm 3.72 [-8.9, 7.41]$ | 0.84 | | |
| Sleep Variability 6 Mo | $0.37 \pm 0.26 [-0.2, 0.94]$ | 0.18 | | |

